# Targeted IS-element sequencing uncovers transposition dynamics during selective pressure in enterococci

**DOI:** 10.1101/2022.08.24.505136

**Authors:** Joshua M. Kirsch, Shannon Ely, Madison E. Stellfox, Karthik Hullahalli, Phat Luong, Kelli L. Palmer, Daria Van Tyne, Breck A. Duerkop

## Abstract

Insertion sequences (IS) are simple transposons implicated in the genome evolution of diverse pathogenic bacterial species. Enterococci have emerged as important human intestinal pathogens with newly adapted virulence potential and antibiotic resistance. These genetic features arose in tandem with large-scale genome evolution mediated by mobile elements. Pathoadaptation in enterococci is thought to be mediated in part by the IS element IS256 through gene inactivation and recombination events. However, the regulation of IS256 and the mechanisms controlling its activation are not well understood. Here, we adapt an IS256-specfic deep sequencing method to describe how chronic lytic phage infection drives widespread diversification of IS256 in *E. faecalis* and how antibiotic exposure is associated with IS256 diversification in both *E. faecalis* and *E. faecium* during a clinical human infection. We show through comparative genomics that IS256 is primarily found in hospital-adapted enterococcal isolates. Analyses of IS256 transposase gene levels reveal that IS256 mobility is regulated at the transcriptional level by multiple mechanisms in *E. faecalis*, indicating tight control of IS256 activation in the absence of selective pressure. Our findings reveal that stressors such as phages and antibiotic exposure drives rapid genome-scale transposition in the enterococci. IS256 diversification can therefore explain how selective pressures mediate evolution of the enterococcal genome, ultimately leading to the emergence of dominant nosocomial lineages that threaten human health.

**Author Summary:** Insertion sequence (IS) elements are simple transposons that are ubiquitous in bacteria. In the enterococci, which includes medically relevant species such as *Enterococcus faecalis* and *Enterococcus faecium*, the IS element IS256 is widespread and has been implicated in pathogenesis and antibiotic resistance. Despite the importance of IS256 to the biology of the enterococci, we know very about how this element is regulated and diversifies enterococcal genomes. Here, we show that IS256 is preferentially found in hospital-adapted and virulent strains of the enterococci. In *E. faecalis* V583, a vancomycin resistant blood isolate, IS256 is regulated by multiple transcriptional mechanisms. To understand how IS256 is mobilized, we adapted an Illumina-based deep sequencing method called IS-Seq to find novel IS256 insertions when applying the selective pressure of bacteriophage (phage) predation. Using this method, we found that chronic phage infection drives IS256 diversification of the *E. faecalis* V583 genome. Additionally, we tracked IS256 insertional activity during a human *E. faecium* infection and found that increased IS256 diversity was associated with specific antibiotic usage. Together, our results demonstrate that the enterococci control IS256 activity to diversify their genomes which may lead to the emergence of hospital-adapted strains that threaten human health.

## Introduction

Enterococci, including the human commensals *Enterococcus faecalis* and *Enterococcus faecium*, are opportunistic pathogens of public health concern due to their acquisition of antibiotic resistance and virulence traits[1]. Enterococcal infections are a leading cause of infective endocarditis [2, 3] and 30% of nosocomial infections are vancomycin resistant [4, 5]. Antibiotic resistance in enterococci is often encoded on mobile genetic elements, including plasmids and composite transposons [6]. Plasmids, especially those containing Inc18, Rep_3, and RepA-N replicon-containing plasmids, frequently encode resistance genes to diverse antibiotics, including chloramphenicol, erythromycin, and gentamicin [7, 8]. Composite transposons are typically organized with a cargo gene (such as an antibiotic resistance gene) flanked by insertion sequence (IS) elements. Examples in the enterococci include Tn4001 or Tn4031, which carries gentamicin resistance genes [9] and Tn1546 and Tn1547, which carry the vancomycin resistance *vanA* and *vanB* operons, respectively [10, 11].

To combat clinically relevant enterococcal strains, new therapeutics are urgently needed. Recently, the concept of using bacterial viruses (bacteriophages or phages) to treat multi-drug resistant (MDR) bacteria is being revisited [12]. Phage therapy case studies show that these viruses can be efficacious against refractory MDR bacteria in human patients [13–17]. However, bacterial resistance to phage infection develops quickly [18–21] and the lessons learned from the past century of antibiotic use suggest that bacteria will gain resistance to phage infection after widespread phage treatment [22]. Acquired phage resistance, although beneficial to bacteria in the face of phage pressure, can come with fitness costs that dampen antibiotic resistance and virulence [12, 23–25]. Therefore, it is imperative to fully understand the mechanisms that promote phage resistance in bacteria, and the phenotypic outcomes of phage resistance. With this knowledge it will be possible to use phage resistance as a tool that can be leveraged against bacteria to enhance current antibacterial therapeutics.

Enterococci have remarkably diverse genomes containing numerous mobile elements that contribute to their adaptation and evolution [26–28]. These include plasmids, prophages, and transposable insertion sequence (IS) elements. Pathogenic enterococci frequently contain multiple IS elements, and IS elements appear to be a feature of newly-adapted nosocomial strains [29]. In the widely studied nosocomial type-strain *E. faecalis* V583, there are 38 IS elements consisting of 11 different types [26]. IS256 is the most abundant IS element in the *E. faecalis* V583 genome, with six chromosomal copies and four plasmid copies. IS256 is also common in other gram-positive bacteria, including staphylococci, where it has been widely studied [30–32]. In the staphylococci, insertion of IS256 into different genes causes clinically-relevant phenotypes, such as small colony variation [33], biofilm formation [31], and antibiotic resistance [34]. The transcription factor σ-B negatively regulates IS256 transposition in the staphylococci through the production of a 3’ antisense RNA [31, 35]. IS256 is a crucial component for enterococcal genome adaptation. In *E. faecalis* V583, the virulence factor cytolysin is attenuated by IS256 and related IS905 insertions [36]. Additionally, copies of IS256 in the *E. faecalis* V583 pheromone-responsive conjugative plasmids pTEF1 and pTEF2 recombine with chromosomal IS256 copies to mobilize broad regions of the *E. faecalis* genome [37]. A related IS element, IS16, has been shown to be highly abundant in hospital-adapted *E. faecium* isolates, suggesting that IS16 may aid in the success of *E. faecium* as a nosocomial pathogen [38]. Considering IS elements are likely involved in the pathoadaptation of the enterococci, little is known about how these elements are regulated and what events lead to their activation.

In this work, we investigated both the regulation and dynamics of IS256 transposition in *E. faecalis* and *E. faecium*. We found that IS256 is present in multiple, genetically disparate isolates of both species, and is enriched in hospital adapted lineages. At steady state, IS256 produces a wealth of low abundance insertions throughout the chromosome and is regulated at the transcriptional levels to tightly control activation. We discovered that phage infection and antibiotic usage in different biological settings were associated with increased diversity of IS256 insertions. In the case of phage infection, cell populations with diversified IS256 insertions maintained phage genomes for an extended period of time and a subpopulation of these cells chronically shed phage particles throughout growth. Phage shedding provided a competitive advantage during co-culture with phage-susceptible enterococci, but this advantage is lost when co-cultured with phage-resistant enterococci. This suggests that phage genome carriage is a strong selective pressure that drives IS256 diversification and can be used to occupy an environmental niche when competing with phage-susceptible bacterial strains. Lastly, we investigated how IS256 diversifies enterococcal genomes in a chronically infected human, and found that IS256 insertion abundances in vivo coincided with specific antibiotic usage. Together, this work sheds light on how IS elements are regulated and expanded in enterococcal genomes during both phage predation and clinical infection, and provides evidence for phage carriage as an important selective pressure that promotes IS256 diversification. Furthermore, this work suggests that therapeutic use of both phages and antibiotics could cause rapid and widespread enterococcal genome evolution for which the physiological consequences are unknown.

## Results

### IS256 is common within enterococcal lineages associated with hospital adaptation and pathogenesis

IS256 is an important factor driving the pathoadaptation of *E. faecalis* [36], yet it is unknown how widely IS256 is distributed among *E. faecalis* strains. To answer this, we searched all available *E. faecalis* genomes from NCBI RefSeq (2065 genomes) for IS256 transposases with 100% amino acid similarity to the IS256 transposase copies found in *E. faecalis* V583. A total of 232 genomes contained one or more IS256 sequences (Fig. 1A). The sporadic distribution of IS256 within the *E. faecalis* phylogenetic tree suggests that IS256 is a recent addition to many *E. faecalis* genomes and has arisen within multiple discrete lineages of this species. To further characterize the distribution of IS256 across *E. faecalis* strains, we performed *in silico* multilocus sequence typing (MLST) on all *E. faecalis* genomes and compared the proportions of genomes with and without IS256 in each sequence type (ST) (Fig. 1B). IS256 occurs within a variety of ST clades and is most abundant in in ST6, ST103, ST778, and ST388 genomes, all of which are associated with hospital-adapted, opportunistic pathogenic lineages (Table S1A) [39–42]. We compared the abundances of different virulence factors between genomes with and without IS256 (Fig. 1C & Table S1C). IS256-containing genomes are enriched for multiple virulence factors compared to genomes lacking IS256, demonstrating that IS256 preferentially occurs in hospital-acquired and virulent *E. faecalis* strains. We next expanded this analysis to include all *E. faecium* genomes from RefSeq (2306 genomes) and similar to *E. faecalis*, *E. faecium* IS256 is primarily found in the nosocomial lineages ST17, ST664, ST736, and ST18 (Fig. 1D-E and Table S1B) [43, 44]. Although IS256 is significantly enriched in virulent *E. faecium* genomes, this was less than *E. faecalis* (Fig. 1F and Table S1D). Finally, IS256 is more frequently found in *E. faecium* genomes compared to *E. faecalis* genomes, confirming prior research (Fig. 1G) [45]. Together these data show that IS256 is widely distributed within the enterococci and is preferentially found in nosocomial and virulent isolates.

**Figure 1.**
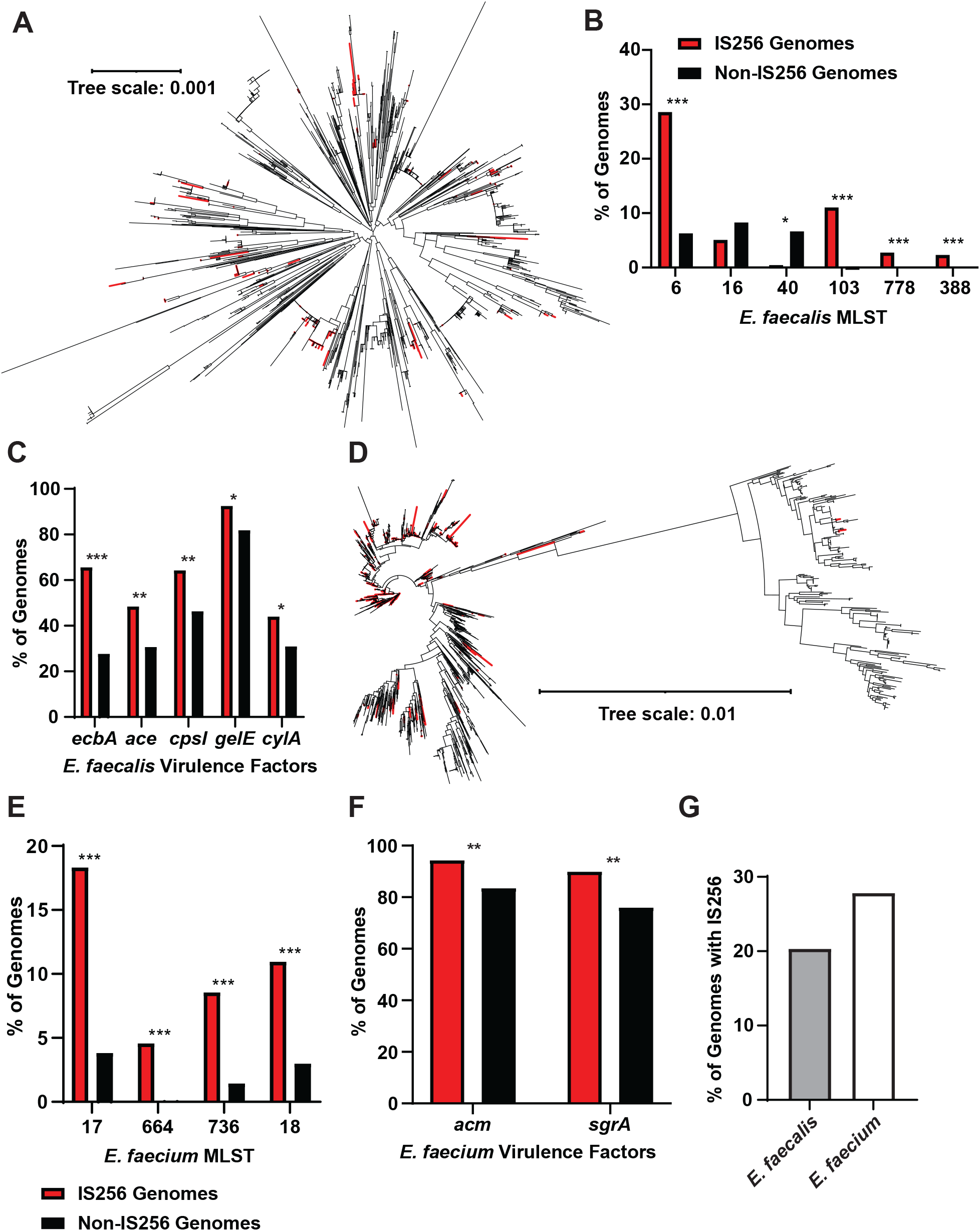
Distribution of IS256 in *E. faecalis* and *E. faecium.* **A)** Phylogenetic tree of *E. faecalis* genomes. Genomes with IS256 are labeled in red and genomes without IS256 are labeled in black. **B)** Presence or absence of IS256 in the genomes of hospital adapted *E. faecalis* MLST types. **C)** Comparison of *E. faecalis* virulence factor abundances in the genomes of hospital adapted *E. faecalis* MLST types. **D)** Phylogenetic tree of *E. faecium genomes*. Genomes with IS256 are labeled in red and genomes without IS256 are labeled in black. **E)** Presence or absence of IS256 in the genomes of hospital adapted *E. faecium* MLST types. **F)** Comparison of *E. faecium* virulence factor abundances in the genomes of hospital adapted *E. faecium* MLST types. **G)** Percent of *E. faecalis* and *E. faecium* genomes from RefSeq that contain IS256 (* *p*<0.05, ** *p*<10^−5^, *** *p*<10^−25^, determined with chi-squared test).

### IS-Seq identifies widespread movement of IS256 in E. faecalis that is transcriptionally and translationally controlled

Previously, IS256 sequences were identified as hot-spots in *E. faecalis* V583 that facilitated the integration of the endogenous plasmids pTEF1 and pTEF2, leading to the mobilization of the pathogenicity island and other chromosomal regions [37]. Instances of IS256 insertions leading to the inactivation of *E. faecalis* genes involved in diverse phenotypes have been described [19, 46–48]. However, we lack a complete understanding of IS256 mobility across the *E. faecalis* genome during selective pressures that would potentially drive IS256 activation and genome diversification. To assess genome-wide IS256 insertions in *E. faecalis* we adapted a next-generation sequencing (NGS) enrichment technique for use with IS256. This technique, IS-Seq, includes IS256 amplicon enrichment during NGS library construction followed by read mapping of IS256-chromosomal junctions to identify specific IS256 insertion locations (Fig. S1) [49]. Similar techniques have been used to identify IS element insertions in *Acinetobacter* [50], and *Mycobacterium* [51]. IS-Seq of wild-type (WT) *E. faecalis* V583 under steady state conditions identified the known IS256 locations in the chromosome and in the three pTEF plasmids (Fig. 2A & S2). One of the IS256 copies on pTEF1 lacks a canonical left inverted repeat (IR), preventing binning of the IS-Seq reads originating from this locus. Numerous low abundance IS256 insertions are observed throughout the *E. faecalis* genome, indicating promiscuous movement of the element within subpopulations of cells (Fig. 2A). The three biological replicate cultures tested in Fig. 2A show that each *E. faecalis* population analyzed has a unique repertoire of IS256 insertions. We hypothesize that these low abundance steady-state insertion events provide a mechanism for rapid genome diversification following exposure to selective pressures.

**Figure 2.**
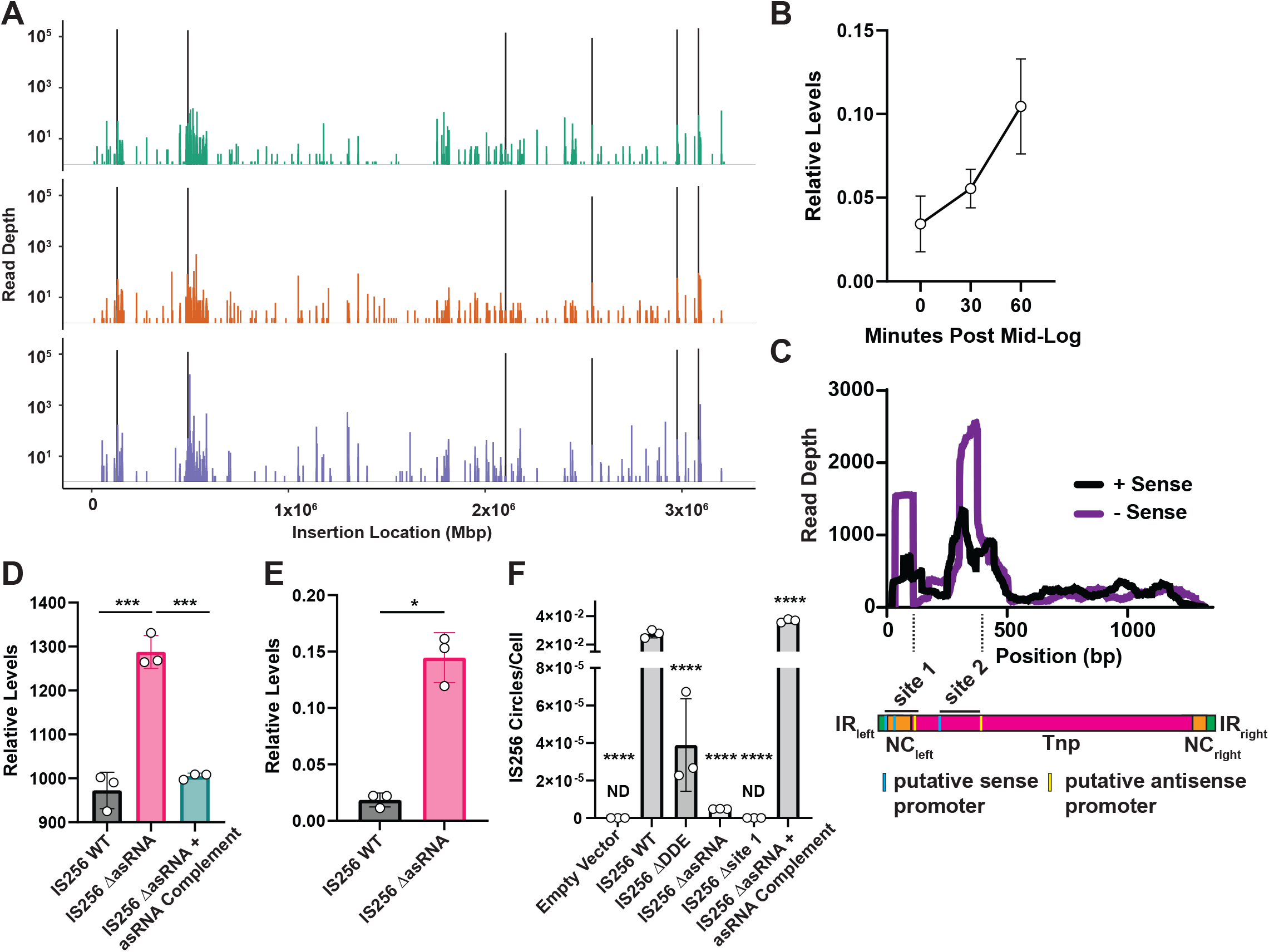
Regulation of IS256 Tnp activity. A) IS256 sequencing (IS-Seq) from three separate biological replicates of *E. faecalis* V583. Peaks in black represent the six endogenous IS256 insertions identified in the V583 chromosome by whole genome sequencing. **B)** RT-qPCR of IS256 *tnp* gene levels relative to the housekeeping ribosomal gene 16s (three biological replicates performed in triplicate). **C)** Stranded RNA-Seq of IS256 tnp gene levels in *E. faecalis* V583 (three biological replicates). **D)** IS256 Tnp protein levels indicated by measuring GFP fluorescence of an IS256-GFP and IS256-GFP ΔasRNA translational fusions. (** *p*<0.01, determined with t-test from three biological replicates). **E**) RT-qPCR of IS256 Tnp RNA levels of WT IS256-GFP and IS256-GFP ΔasRNA (* *p*<0.05, determined with t-test from three biological replicates). **F**) qPCR of IS256 circular intermediates in *E. faecalis* OG1RF carrying either an empty vector (EV), IS256, catalytically inactive IS256, IS256 ΔasRNA, or IS256 site 2 Only (**** *p*<0.0001, determined with one-way ANOVA. Every treatment was compared to the IS256 treatment and corrected for multiple comparisons using the Dunnett method).

To understand how IS256 is regulated in *E. faecalis* under steady state conditions, we investigated possible mechanisms of repression of the IS element. First, we assessed the transcription of the IS256 transposase (*tnp*) gene. During growth in rich media, IS256 gene transcription modestly increases during exponential growth, and reaches 1/10^th^ of the levels of the reference 16S rRNA gene (Fig. 2B). Considering there are 10 copies of IS256 and 4 copies of 16S in *E. faecalis* V583, each IS256 copy achieves 1/25^th^ the transcriptional activity of a single 16S gene if all copies are expressed similarly. Thus, the relatively weak IS256 promoter may limit over-activation of the element. Next, we investigated whether IS256 is further transcriptionally or translationally regulated. Other IS elements have been reported to be controlled by antisense small RNAs (asRNA) [52]. These asRNAs repress transposase translation through diverse mechanisms, including binding to sense transcripts at or before the start codon and ribosome binding site to prevent translation of the sense transcript or targeting the *tnp* transcript for RNase III cleavage [53]. We reanalyzed a publicly available stranded RNA-Seq dataset of *E. faecalis* V583 [54] by aligning reads to an IS256 element with delineation of the sense and antisense alignments (Fig. 2C). The transcriptional start site (TSS) of IS256 has not been experimentally confirmed in any bacterial species and is predicted to reside in the 5’ terminus of the element [55]. We demonstrate that IS256 has two TSSs indicated by high sense read mapping coverage in distinct peaks. Additionally, each sense peak is overlayed by a higher antisense peak, demonstrating that antisense inhibition is present at both TSSs. If translation initiates at the site-2 TSS this would produce a truncated IS256 Tnp, here after referred to as IS256 Tnp site 2. Upon further investigation of these TSSs and their asRNAs, we found that the site-1 asRNA is likely under the control of a canonical σ-70 promoter. To identify if this asRNA controls IS256 Tnp translation, we constructed an IS256 Tnp-GFP translational reporter. This construct consists of an IS256 Tnp coding sequence fused in frame at its C-terminus to a green fluorescence protein (GFP) gene. GFP fluorescence serves as a proxy for translation of the IS256 Tnp. In addition, we built a version of the IS256 Tnp-GFP fusion lacking the predicted −10 element of the site-1 asRNA. We transformed these constructs into *E. faecalis* OG1RF, a strain that lacks native IS256 elements and IS256-derived asRNAs in its genome, and measured GFP fluorescence in these cells (Fig. 2D). In cells lacking the site-1 asRNA, there is significantly higher GFP fluorescence compared to cells with an intact site-1 asRNA promoter, demonstrating that these asRNAs repress IS256 Tnp translation. This phenotype was fully complemented. We also measured the RNA levels of the IS256 *tnp* gene from the same constructs and found that there were significantly more *tnp* transcripts in the IS256-GFP ΔasRNA cells (Fig. 2E). These results show that in *E. faecalis* IS256 is regulated by multiple mechanisms at the transcriptional level.

To extend our findings, we compared the abundance of IS256 circular intermediates between different IS256 constructs in *E. faecalis* OG1RF (Fig. 2F), a strain which lacks any IS256 elements. IS256 elements circularize during transposition, which can be used to measure active transposition [30, 31]. Using a multi-copy plasmid backbone, we compared the rate of IS256 circularization in *E. faecalis* OG1RF carrying the following constructs: WT IS256, an inactive IS256 gene lacking the coding sequences for the two catalytic aspartic acid residues of the Tnp’s DDE motif (ΔDDE), an IS256 lacking the promoter for the site 1 asRNA (IS256 ΔasRNA), and an IS256 lacking the 5’ UTR and coding sequences upstream of the site 2 promoter (IS256 site 2). The IS256 site 2 construct retains the 5’ IR. Diagrams of these constructs are shown in Fig. S3. Surprisingly, we found that all constructs produced significantly less IS256 circles compared to WT IS256. This suggests that the site 2 Tnp product and IS256 ΔasRNA are inactive for transposition, even though loss of the asRNA increases IS256 Tnp transcription. To confirm these findings, we complemented the asRNA and found that this restored the phenotype and increased IS256 circular intermediates above WT IS256 levels. Additionally, we confirmed that IS256 circles require an active IS256 element as there were significantly less IS256 circles produced by the inactive IS256 element.

### IS256 selection during phage predation

Our previous work suggested that phage predation increases IS256 transposition in *E. faecalis* V583 and is used as a mechanism of phage resistance [18]. To determine if phage selective pressure results in the diversification of IS256 elements in *E. faecalis*, we challenged *E. faecalis* V583 with the phage phi19 [18]. Following phi19 exposure, we isolated single colonies that were resistant to phi19. Whole genome sequencing (WGS) of these phage-resistant isolates predicted new IS256 insertions in phi19 resistant isolates (referred to as 19RS strains) (Fig. 3A). Among the new IS256 insertions in 19RS strains, we found an insertion in the 3’ end of *epaX*, a glycosyltransferase gene involved in the biosynthesis of the enterococcal polysaccharide antigen that is required for infection by phi19 [18]. To verify this insertion and its orientation, we performed PCR using genomic DNA (gDNA) from four 19RS strains and WT *E. faecalis* V583 (Fig. 3B). We found that all 19RS strains harbored an IS256 insertion in *epaX* in both the forward and reverse complement orientations. This IS256 insertion likely prevents functional EpaX from being expressed blocking phage infection. Recently, Lossouarn et al. identified IS256 insertions inactivating *epaX*, confirming that an IS256 insertion in *epaX* is sufficient to prevent phage infection [48]. These results demonstrate that 19RS strains contain a novel IS256 insertion not found in WT *E. faecalis* V583. These results also suggest that 19RS strains, even though they were isolated as individual colonies, are heterogeneous populations with varied abundances of IS256 insertions.

**Figure 3.**
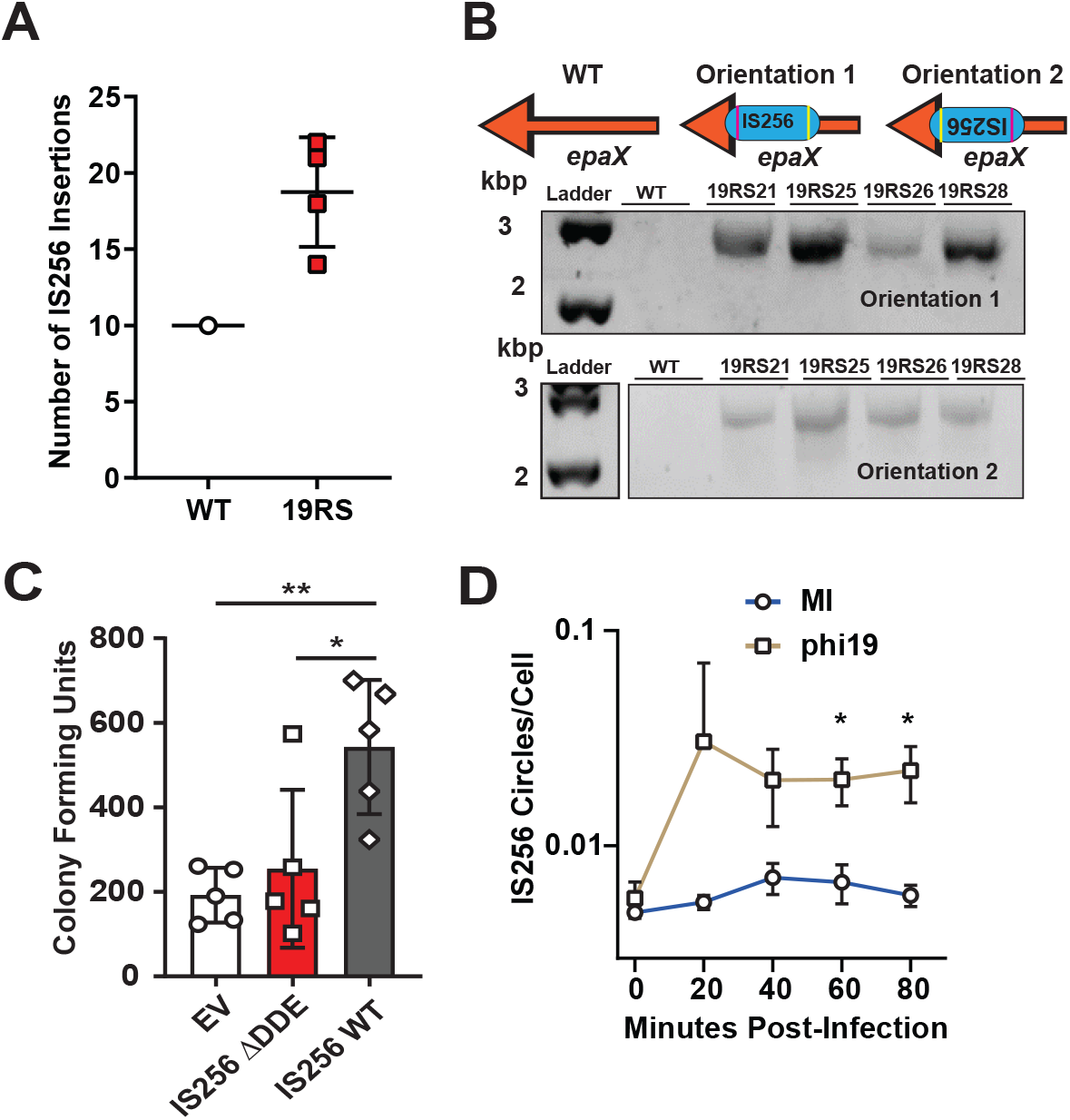
IS256 activity confers phage resistance and is elevated in *E. faecalis* isolates resistant to phi19. A) Enumeration of predicted IS256 insertions determined by whole genome sequencing of a single wild type *E. faecalis* V583 isolate and four independent phage resistant 19RS strains. A value of 10 indicates the total number of IS256 copies found in the *E. faecalis* V583 reference genome (GCA_000007785.1). **B)** Representative PCR showing that IS256 insertions occur in both the positive and negative strand orientations in the *epaX* gene 19RS strains. **C)** Emergence of phage resistant *E. faecalis* OG1RF with IS256 or a catalytically dead (ΔDDE) IS256. EV indicates *E. faecalis* OG1RF carrying the empty vector. (* *p*<0.05, ** *p*<0.01, determined by multiple comparison t-test using five biological replicates). **D)** qPCR quantification of IS256 circular intermediates normalized to the single copy gene *clpX* following phi19 infection or mock infection (MI). (* *p*<0.05, determined with multiple comparison t-test from three biological replicates performed in triplicate).

To test if IS256 provides an advantage to *E. faecalis* against phage infection, we used phi19 to infect *E. faecalis* OG1RF carrying either an empty vector (EV), WT IS256, or the inactive version of IS256 (ΔDDE) (Fig. 3C). We found that *E. faecalis* cells containing a functional IS256 gene were more resistant to phage infection compared to cells containing the catalytically dead Tnp or the empty vector, indicating that IS256 carriage increases mutation frequency. To understand if IS256 activation was initiated by phi19 infection, we quantified the number of IS256 circular intermediates. During the course of phi19 infection, IS256 circular intermediates significantly increased relative to a mock-infected cells (Fig. 3D).

WGS analysis indicates that phage predation increases IS256 across the *E. faecalis* genome. However, this method is low resolution. To quantify the exact locations and abundances of IS256 insertions following phage pressure, we performed IS-Seq on 19RS strains and compared these to WT *E. faecalis* V583 (Fig. 4A). We expected that each strain would have substantial IS256 insertional variability, given that 19RS strains contained an IS256 insertion in multiple orientations (Fig. 3B). To provide a rigorous and reproducible assessment of IS256 insertions in these strains, we performed IS-Seq on gDNA collected from three biological replicates for both WT *E. faecalis* V583 and 19RS strains. The three replicates were used to calculate the average peak intensity at each insertion site. IS-Seq revealed a genome-wide diversification of IS256 insertions at new locations in 19RS strains (Fig. 4A & Table S2). Pairwise comparisons determined that 19RS strains are significantly enriched in the number of novel IS256 insertions compared WT *E. faecalis* V583. These comparisons were determined using the DESeq2 [56], which accounts for over-dispersion and provides a rigorous multiple testing statistic. We used raw sequencing read counts of each insertion for this analysis (Fig. 4B). The results of this analysis are detailed in Tables S2 and S3. New IS256 insertions in the pTEF plasmids were also found in the 19RS strains (Fig. S4A-C). Additionally, we measured IS256 circular intermediates in our IS-Seq dataset by aligning IS256 reads (which originate from the left terminus of IS256) to the right terminus of an IS256 element. We found that 19RS strains have a significantly greater number of IS256 circular intermediates compared to WT *E. faecalis* V583 (Fig. 4C).

**Figure 4.**
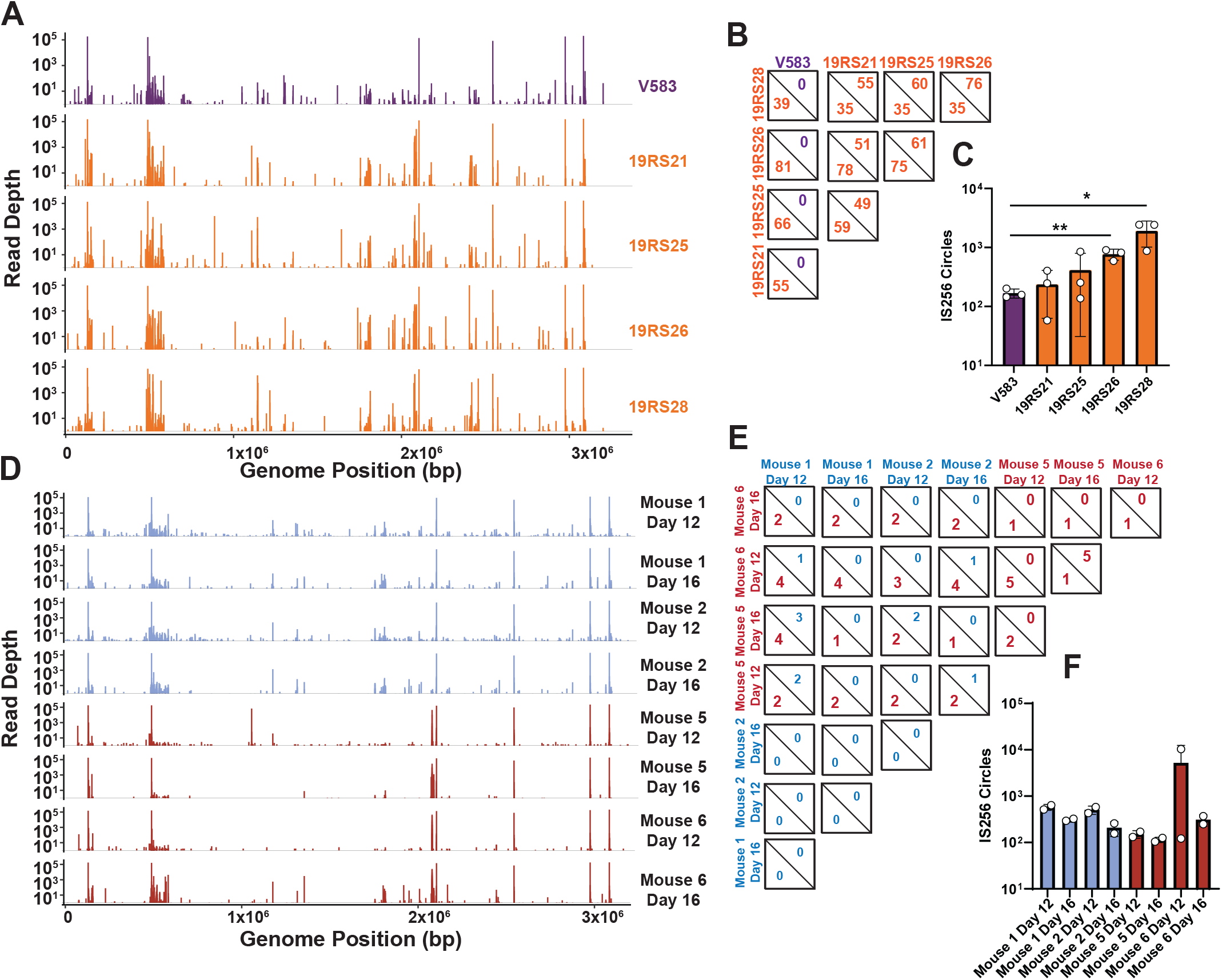
IS-Seq to identify IS256 insertion sites in *E. faecalis* phage resistant isolates in vitro and in vivo. A) IS-Seq of *E. faecalis* V583, and 19RS strains. Each peak in this panel is an average of three biological replicates per strain. **B)** Quantification of significantly enriched insertions between pairwise comparisons of each group in A. In each square, the top value is the number of insertions overabundant in the column group and the bottom value is number of insertions overabundant in the row group. **C)** IS256 circle formation determined with IS-Seq from samples shown in A. (* *p*<0.05, ** *p*<0.01, determined with multiple comparison t-test using three biological replicates). **D)** IS-Seq of *E. faecalis* V583 and phage resistant colonies isolated from the feces of mice colonized with (Mouse 5 & 6) and without (Mouse 1 & 2) oral phi19 inoculation. Each peak in this figure is an average of two biological replicates. **E)** Quantification of significantly enriched insertions between pairwise comparisons of each group in D. In each square, the top value is the number of insertions overabundant in the column group and the bottom value is number of insertions overabundant in the row group. **F)** IS256 circle formation determined with IS-Seq from samples shown in D. Insertions considered significant in panels A and D have an adjusted *p* < 0.05, a fold change > 4, and were supported by at least 100 reads in one group during pairwise comparison. Statistics were computed using the DESeq2 generalized linear model algorithm.

We focused on two genomic regions that experienced a high level of IS256 insertions in the 19RS strains. The first was the *epa* locus. In addition to numerous novel insertions in *epaX*, we found insertions within other *epa* genes including *epaR*, *epaS*, *epaAA* and *epaAB*. (Fig. S5A). IS256 insertions in these *epa* genes would alter the teichoic acid decoration of the core rhammnose-containing Epa, likely altering the surface chemistry of *E. faecalis* resulting in phage resistance [18]. We also identified multiple insertions in the *vex/vnc* operon (Fig. S5B), which is purported to have both virulence and antibiotic resistance functions [57, 58].

*E. faecalis* is a native member of the intestinal microbiota and can outgrow and cause disease during intestinal dysbiosis. Phages have been proposed as a treatment option for difficult to control enterococcal intestinal blooms [59]. To determine if phage predation supported IS256 diversification in the intestine, we colonized mice with *E. faecalis* V583 and orally challenged these animals with phi19 (Fig. S6). We performed IS-Seq on fecal gDNA from these animals and found that phi19 exposure in the intestine also increased new IS256 insertions, albeit to a lesser extent than in vitro (Fig. 4D-F, Fig. S4D-F, & Table S3). We found that similar to cells exposed to phi19 in vitro, IS256 insertions within the same location in *epaX* and the *vex*/*vnc* operons are observed in the mouse intestine following phage administration (Fig. S5C-D). This suggests that phage selection plays a role in IS256 diversification in the context of the intestine.

### Enterococci with expanded IS256 genomes chronically shed phi19 phages

Phage genomes can be maintained in their host bacterium in a variety of configurations, including chromosomally integrated or as cytosolic episomes [60]. In phage-bacteria communities, sensitive hosts, resistant hosts, and hosts containing phage genomes coexist at a consistent ratio [60]. Sensitive hosts are infected with phage particles and either perish, increasing the number of phage particles, become resistant, or become lysogenized. Over time, a subset of lysogenized and/or resistant bacteria desensitize to the phage infection and the cycle repeats. This phenomenon is termed carrier state [60]. Phage genomes can also be in an intermediate state between lytic replication and lysogenic conversion referred to as pseudolysogeny [60]. Such phages can enter the lytic cycle spontaneously or following cell stress, and reports indicate that lysogenic phages can be continuously shed from cells with little to no reduction in viability of the overall bacterial population [61, 62]. During the analysis of bacterial colonies originating from *E. faecalis* 19RS strains, we discovered that the phi19 genome is maintained in these phage-resistant bacteria and the population can grow at a stable rate. This carriage of the phi19 genome may explain its ability to influence IS256 transposition. We found that 19RS strains continually release infectious phage particles during different stages of growth (Fig. 5A). These particles were confirmed to be phi19 using PCR (Fig. 5B). We discovered that ∼10% of 19RS colonies produced infectious phages (Fig. 5C). This suggests that a minority of each population exerts a selective pressure to maintain phage resistance by producing infectious phage progeny. Loss of phage resistance in this population would lead to phage infection and population collapse. Colonies that produced zones of clearing were isolated and serially passaged. Cells that initially produced zones of clearing continued to produce zones for at least three passages (Fig. S7), demonstrating that phi19 is stably carried in this subpopulation. Next, we reanalyzed reads from our WGS experiments and found that we could recover the phi19 genome with high coverage from 19RS strains (Fig. 5D). WT *E. faecalis* V583 did not have any detectable phi19 genomic DNA. To determine if the phage genome was integrated into the *E. faecalis* chromosome or one of its three endogenous plasmids, we performed long-read Oxford Nanopore sequencing on the 19RS strains. This technique provides reads up to 20kb and can be used to find large genomic insertions and rearrangements. We first searched for reads with phage genomic DNA content and then identified if any of these reads contained host genomic DNA. Every 19RS strain had reads containing phage and host hybrid genomic reads. This ranged from 0.2-10% of total phi19 reads (Fig. 5E). While this may be interpreted evidence of phi19 integration, Nanopore reads have a low rate of chimeric read formation. Chimeric reads are reads containing two sequences combined as a result of a technical artifact during library preparation. Chimeric read formation in Nanopore libraries is reported as 1.7% [63] of total reads, which is in agreement with the proportion of phi19 reads containing both phage and host DNA. Thus, we believe that the phage genome is not integrated into the host genome in 19RS cells but is otherwise maintained as an independent genome that could be considered temperate. Collectively, these results show that 19RS strains release phages from a subpopulation of cells, which may impose stress on *E. faecalis* influencing IS256 mobility.

**Figure 5.**
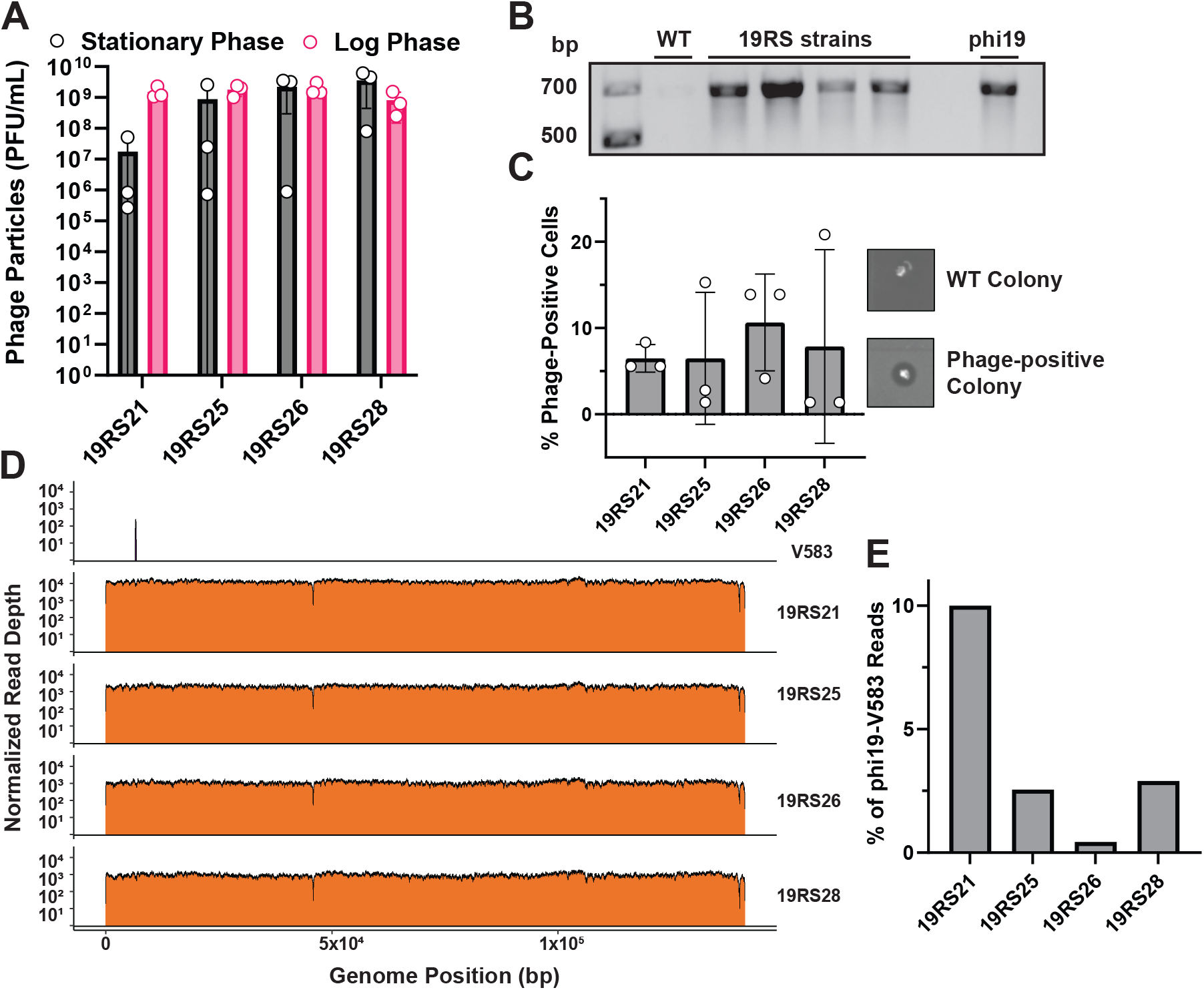
*E. faecalis* 19RS strains chronically shed phi19 virus and harbor the phage genome intracellularly. A) Quantification of shed infectious phi19 phage particles. **B)** PCR confirmation of phi19 recovered from culture fluid of *E. faecalis* 19RS strains. **C)** Proportion of cells actively replicating and releasing infectious phi19 phage particles. **D)** WGS read mapping alignments to the phi19 genome from *E. faecalis* V583 and 19RS strains. Data represent three biological replicates. **E)** Percentage of phi19 nanopore reads from the 19RS strains that contain host sequences.

Considering that 19RS strains have IS256-rich genomes and release infectious phages, we hypothesized that phage carriage may increase the competitiveness of 19RS strains when co-cultured with phage sensitive strains. We performed competition assays where WT *E. faecalis* or 19RS strains were competed against *E. faecalis* OG1RF or an OG1RF Δ*epaOX* mutant strain, without the addition of exogenous phages to the cultures (Fig. 6A-B). The *E. faecalis* OG1RF *epaOX* gene is homologous to *E. faecalis* V583 *epaX* [64] and *E. faecalis* OG1RF Δ*epaOX* is resistant to phi19 infection. 19RS strains that were shedding phi19 outcompeted *E. faecalis* OG1RF. However, they were unable to outcompete *E. faecalis* OG1RF Δ*epaOX*. These results demonstrate that phi19 carriage provides a selective advantage to *E. faecalis* cells when competing with related bacteria that are phage susceptible but if competitors are phage resistant, phi19 carriage comes with a significant fitness cost. To better understand the fitness cost of phage carriage, we compared the growth rate of E. faecalis 19RS strains, V583, and OG1RF in monoculture. *E. faecalis* 19RS strains have a growth defect compared to V583 and OG1RF when measuring optical density (Fig. 6C) and doubling time (Fig. 6D).

**Figure 6.**
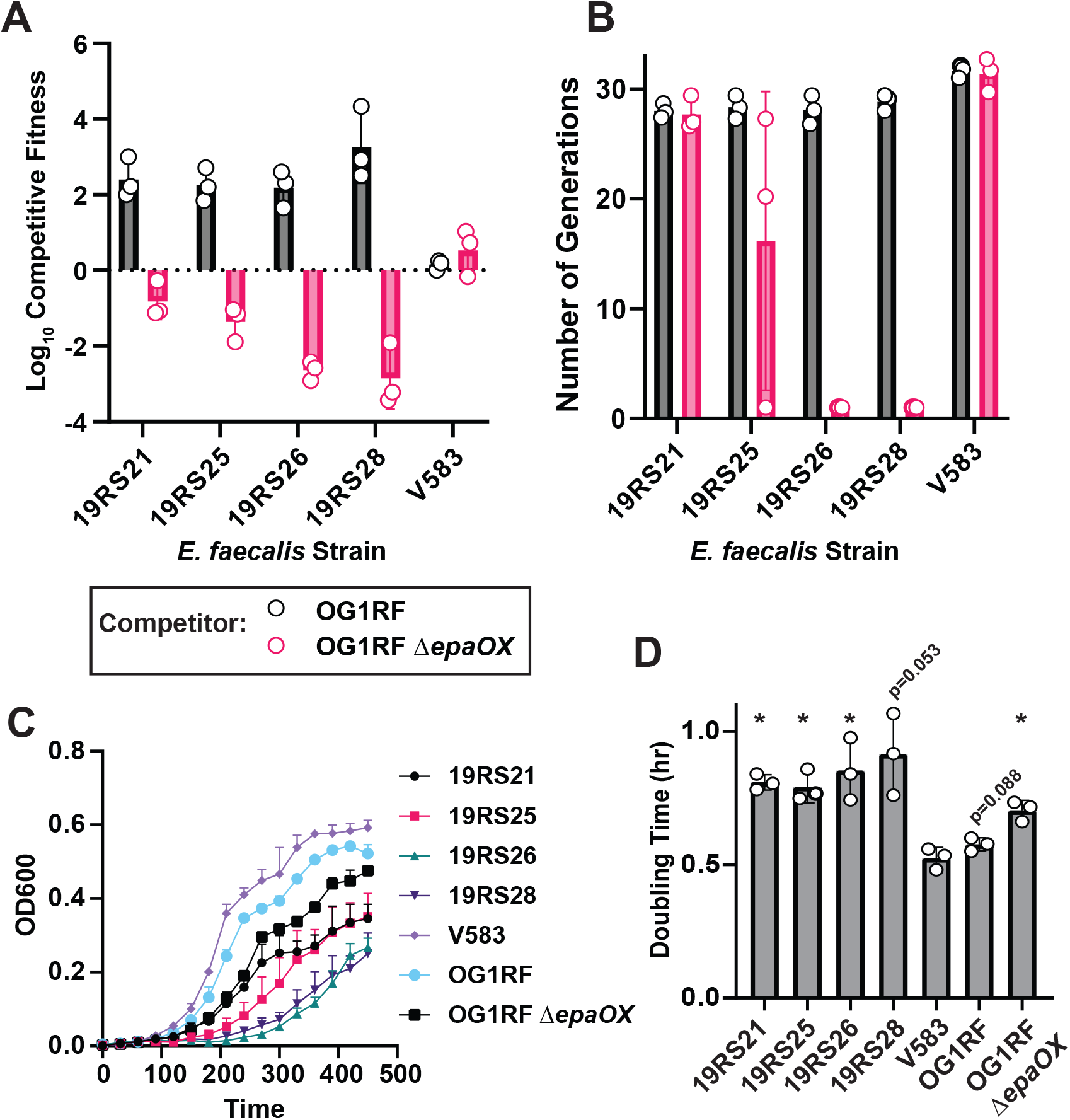
phi19 shedding *E. faecalis* 19RS strains are more fit during competition with a phage susceptible *E. faecalis* strain. A) Competitive fitness of *E. faecalis* V583 and 19RS strains relative to *E. faecalis* OG1RF or an *E. faecalis* OG1RF derivative with a deletion in the *epaX* homolog *epaOX*. Data represent three biological replicates. **B)** Number of generations of V583, 19RS, and OG1RF strains during the competition assay in **A**. **C)** Growth curves of V583, 19RS, and OG1RF strains. **D)** Doubling time in hours of Growth curves of V583, 19RS, and OG1RF strains. (* *p*<0.05, determined with t-tests in comparison to V583 from three biological replicates. Multiple testing correction was performed with FDR method).

### IS256 mobilization occurs in enterococcal genomes during human infection

IS elements contribute to genome diversification and adaptive plasticity that is important for bacterial evolution [65]. Environmental pressure is a strong driver of IS element mobility [66-68], and the work described here expands this knowledge to phage predation. Antibiotics are a strong selective pressure that can promote bacterial evolution, and IS256 insertions have been tied to pathoadaptation in enterococci during human infections [36]. Furthermore, antibiotics can increase IS256 transposition [69] To assess if environmental cues such as antibiotics may also be driving IS element diversification, we investigated whether enterococcal populations within a single individual infected with *E. faecium* experiences IS256 mobility during therapeutic intervention. Stools samples were collected at various timepoints over 109 days, during which the individual was treated with regimens containing daptomycin, vancomycin, and the vancomycin analogue oritavancin for recurrent *E*. *faecium* bacteremia. IS-Seq was performed on these samples across this time series (Fig. 7A). Nearly all of the insertions found using IS-Seq occurred in contigs which were generated from *E. faecium*, rather than *E. faecalis* (Fig. S8). We found that these *E. faecium* populations had a larger proportion of IS256 insertions following oritavancin therapy compared to daptomycin and/or vancomycin therapy. Comparing differentially abundant insertions during oritavancin treatment revealed multiple enriched insertions in the coding sequences of a putative Rib transcriptional regulator, and the *vex*/*vncRS* operon also identified as a IS256 insertion hotspot in *E. faecalis* 19RS strains (Fig. 7B-C). These two operons are associated with antibiotic resistance and virulence in *E. faecium* and IS256 insertions likely render these genes nonfunctional. These insertions increase in sequencing depth during the course of treatment and peak during oritavancin administration. The *vex*/*vncRS* operon in *E. faecalis* V583 contains an IS element belonging to the ISL3 family [26], indicating that the genome evolution of *E. faecium* during blood stream infection may follow a similar trajectory to *E. faecalis* V583 [26]. We also found that IS256 regularly interrupts an aminoglycoside resistance operon (Fig. S9). Oscillating insertions both in coding and non-coding regions in this operon suggest that this region may be an IS256 insertion hotspot. Prophage abundance also increased during oritavancin treatment (Fig. S10). Overabundant prophages had nucleotide sequence homology to the prophage genomes from *E. faecium* strains, but lacked homology to enterococcal prophages known to carry platelet binding protein genes [70] or to enterococcal prophages associated with increased colonization ability [71]. These results suggest that antibiotic treatment can also drive the diversification of multiple classes of mobile elements in the human host. Lastly, we found that oritavancin-treated samples had an increase in IS256 circle formation indicating that in addition to novel insertion events, active IS256 transposition is occurring during blood stream infection (Fig. 7D). Overall, these findings suggest that IS256 transposition is frequent in the intestine during *E. faecium* infection, and this transposition likely impacts the antibiotic resistance and virulence profiles of the infecting bacterial population.

**Figure 7.**
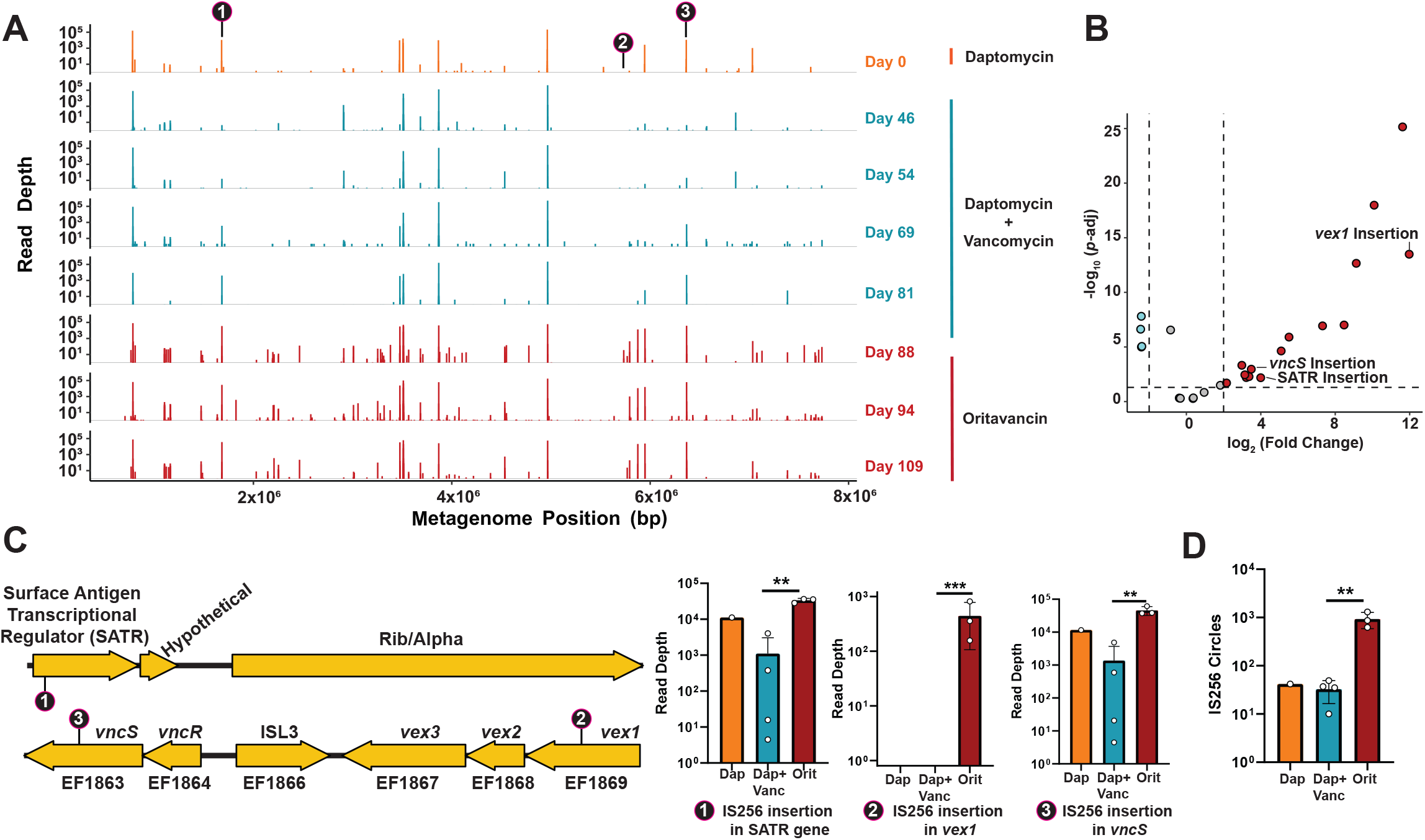
IS-Seq of a longitudinal series of stool samples from a human patient with a vancomycin resistant enterococcal infection. A) Insertions found via IS-Seq plotted on a concatenated “metagenome” of contiguous enterococcal sequences from stool genomic DNA. IS256 sequences were mapped against individual contigs to identify IS256 insertion locations. Then all contigs were concatenated together to build a linearized representation of the metagenome. Each insertion is derived from one biological replicate per time point from the patient’s stool. **B)** Volcano plot of differentially abundant insertions during Daptomycin+Vancomycin co-therapy and Oritavancin therapy. Enriched insertions (dark red) were overabundant during Oritavancin therapy. Significant insertions have an adjusted *p* < 0.05, a fold change > 4, and were supported by at least 100 reads in one group during pairwise comparisons. Statistics were computed using the DESeq2 generalized linear model algorithm. **C)** Gene organization of loci where elevated IS256 insertions occurred are indicated by arrows. Lollipops show the location of IS256 insertion and bar graphs are read depth abundances found for each insertion. (* *p*< 0.05, **, *p*< 0.01, ***, *p*< 0.001, determined with DESeq2) **D)** IS256 circle formation for each treatment group inferred from IS-Seq. (** *p*< 0.01, determined with unpaired t-test using 3-4 biological replicates).

## Discussion

IS elements directly shape bacterial physiological responses involved in antibiotic resistance and virulence, and are key drivers of genome evolution [72–74]. Considering the importance of IS element biology to bacterial fitness, the selective pressures that guide IS element regulation and mobility remain poorly defined. Using the Gram-positive opportunistic pathogen *E. faecalis,* we extend our knowledge of IS elements by detailing the population biology, regulation, and transposition dynamics of IS256. IS256 is a widely distributed IS element in Gram-positive pathogens. Through the use of bacterial genetics, comparative genomics, and IS-Seq we have revealed that IS256 insertion events are strongly dictated by phage predation and the mammalian host environment. Together these results suggest that IS256 mobility creates a genetically flexible genome that enables enterococci to rapidly adapt to diverse environmental conditions.

IS256 is common in *E. faecalis* genomes and is found more frequently in hospital adapted lineages (Fig. 1). Consistent with this idea, IS256 element abundance is a defining feature of hospital adapted *E. faecium* isolates [43, 44]. Movement of IS elements between cells requires other mobile elements, such as phages and plasmids, as IS elements lack any transforming or transducing ability [27]. The pattern of intermittent IS256 abundance in the enterococci is likely due to periodic exposure to IS256-containing mobile elements. In other words, genetically-related strains may interact with IS256-containing mobile elements at different rates, resulting in the dissemination of IS256 to only a subset of the population. The enrichment of IS256 among hospital adapted strains may reflect the need for increased genomic plasticity in these strains as they adapt to an altered host environment [44, 75]. Pathogens can utilize IS elements to form pseudogenes and to condense genome size through IS-mediated recombination, which can inhibit immune detection through loss of virulence factors and reduces the genome’s metabolic burden [74, 75].

Regulation of IS256 has not been investigated in the enterococci and is key to understanding this element’s biology. Here we show that IS256 is basally active in *E. faecalis* V583 and is regulated by multiple mechanisms at the transcriptional level (Fig. 2B-E). This is important because overactive IS256 could be catastrophic to *E. faecalis* viability and layers of regulation likely temper the elements activity until appropriate conditions require IS256 activation. It has been shown previously that transposase protein levels are correlated to transposition frequency in *E. coli* [76, 77]. Additionally, we show that IS256 has two different TSSs as part of this regulation and that both sites are regulated by asRNAs. The protein product occurring from the site-2 TSS is predicted to be truncated. Other IS elements encode two or more protein coding frames and these different protein products can contribute to IS element regulation through inhibition [73, 78]. To better understand this, we built a truncated version of the IS256 Tnp and showed that it is inactive (Fig. 2F). Previous work has shown that similarly truncated IS256 variants are able to bind to the element’s IRs, but structural studies indicate that this truncation likely loses key dimerization domains essential for transposition [79, 80]. We hypothesize that IS256 Tnp site 2 binds to the IRs to prevent full-length IS256 Tnp from binding and activating transposition.

asRNAs are known to regulate transposase protein levels in the *E. coli*, but to our knowledge, these have not yet been investigated in the enterococci. Here, we show that the asRNA produced from site1 in IS256 negatively regulates the RNA levels of the IS256 Tnp. asRNAs can repress transcription through formation of transcriptional repressors or through targeting of the mRNA for degradation by RNAse III [53, 81]. Additionally, loss of this asRNA led to a dramatic decrease in IS256 circular intermediates (Fig. 2F). We hypothesize that this asRNA plays a crucial role in controlling IS256 transposition.

Using IS-Seq as a readout of basal IS256 diversity, we show that IS256 insertions function as a mechanism of genome-wide mutation (Fig. 2A). This is the highest resolution IS-Seq map created, to our knowledge, and demonstrates that this technique can identify low-abundance insertions with high confidence [50, 51]. Baseline IS256 movement likely adds mutational diversity to an enterococcal population which allows for adaption to changing environmental conditions through selection.

Previous work from our group determined that *E. faecalis* infection by the phage phi19 was associated with elevated of IS256 insertions [18] (Fig. 3A). This was found using WGS that was based on less than 10 reads per insertion. We extended this observation by performing IS-Seq to accurately identify these novel insertion sites both in vitro and in vivo. We found that *E. faecalis* challenged with phi19 (19RS strains) had a major increase in new IS256 insertions (Fig. 4A-B). 19RS strains were chronically infected by phi19 and cells within the population carried the phi19 genome and this chronic infection may provide a continuous selective pressure that aids in IS256 mobility. This correlated with 19RS strains harboring elevated IS256 circular intermediates (Fig. 4C). These circles are formed during IS256 copy-paste transposition and have been studied extensively [30].

We also observed that in a mouse model of *E. faecalis* V583 intestinal colonization, exposure to phi19 resulted in a similar diversification of IS256 (Fig. 4D-F). IS256 movement was less robust in the mouse intestine compared to the levels seen in vitro. This may be for several reasons, including a lower frequency of insertion events per location in vivo and a lower number of samples sequenced. An additional explanation is that Epa is crucial for bacterial colonization and *epa* mutations may lead to these mutants being outcompeted in the intestine [18, 82].

Although we found evidence for phage-mediated IS256 activation in *E. faecalis* 19RS strains, we also determined that selection is important for the location of IS256 insertions (Fig. 4G-H). Cells that were not exposed to phi19 had lower IS256 read density in *epaX*. *EpaX* is a glycosyltransferase involved in teichoic acid decoration of the core rhamnose containing Epa exopolysaccharide backbone. Mutations in *epaX* result in poor phage adsorption [18]. We found no other insertions in genes known to be involved in phage infection indicating that *epaX* and other *epa* genes are likely hotspots for IS256 integration following phage pressure. In light of these discoveries, we hypothesize that initial phage infection may select for *epa*-IS256 insertions and that prolonged phage exposure though chronic infection leads to increased and non-specific IS256 transposition as a stress response. *epa* mutations have been shown to sensitize *E. faecalis* to cell wall targeting antibiotics and blood killing and reduces virulence in animal models [18, 83, 84]. This suggests that even though phage carrier state rapidly evolves the *E. faecalis* population, it could also reduce the populations virulence potential. We suspect that phage carriage and release will continually pressure intermittently resistant cells and force fully resistant cells to maintain high levels of resistance. Carrier state is also associated with genetic adaption in the *Bacteroides*, a prevalent member of the gut microbiota. *Bacteroides* coexist with crAss-phages in a constant cycle of infection, resistance, and re-sensitization. To counter crAss-phage, *Bacteroides* will reversibly invert the genetic sequences of capsule genes to prevent phage adsorption and attachment [85].

IS elements are involved in pseudogene formation during pathoadaption. Examples include IS256 insertions in the cytolysin operon of *E. faecalis* V583 [36] and a dependence on IS insertions in *E. coli* to evolve resistance to macrophage killing [86]. Additionally, how enterococci evolve during human infection is not well understood and may involve a variety of genetic changes in cell wall modifying [87] and cellular respiration [88] genes. While IS elements are directly involved in enterococcal pathoadaptation [36], an in-depth analysis of IS-mediated mutations during human infection had never been performed. Previously, such dynamics were studied by laboriously sequencing individual isolates. While this approach can successfully identify new IS element insertion sites, it overlooks IS element mobility within the entirety of the bacterial population. Here, we show that IS-Seq can be used to study bacterial adaptation at the population level and can reveal multiple insertions within such a population (Fig. 7). Following oritavancin therapy an increase in IS256 insertions occurred in the *vex*/*vncRS* operon, which has been implicated in both virulence [58] and vancomycin resistance [57]. Additionally, there is an increase in insertions in a putative transcriptional regulator that may regulate a Rib/Alpha adhesin [89, 90]. Loss of these potential virulence factors may lead to modulation of virulence and avoidance of immune activation or detection. We predict that these insertions arise from an increase in IS256 mobility given the increase in IS256 circular intermediates (Fig. 7D), followed by selection for beneficial insertions. This is likely similar to phage selection for insertions in the *epa* operon.

Additionally, we found highly abundant IS256 insertions in an aminoglycoside acetyltransferase. Vancomycin is reported to act synergistically with gentamicin against enterococci [91], likely due to increased gentamicin transport through the vancomycin-permeabilized cell wall [92]. In recent years, enterococci with aminoglycoside resistance have emerged [93] with acquisition of resistance genes carried in an IS256 composite transposon [94]. Our results suggest that genetic modulation of aminoglycoside resistance fluctuates in the host during infection, with different highly abundant insertions prominent at different timepoints. This data suggests that aminoglycoside resistance is repeatedly modulated and the intestinal enterococcal population regularly samples different versions of this aminoglycoside resistance operon. Future studies to identify if glycopeptide treatment potentiates aminoglycoside sensitivity through IS256 insertions is warranted. We additionally show that oritavancin treatment is associated with IS256 diversification in this patient. Oritavancin is an analog of vancomycin, but is also reported to disrupt cell membrane integrity and possess additional mechanisms for inhibiting cell wall synthesis [95]. This may explain the difference in IS256 expandability, as this antibiotic may impose strong selective pressure resulting in a broader range of mutations. Likewise, oritavancin treatment was associated with an increase in IS256 circles. In *Staphylococcus aureus*, vancomycin treatment increases IS256 transposition [35, 69], reflecting a similar glycopeptide-induced IS256 activation.

In summary, we show how the widespread and clinically-relevant IS element IS256 creates genome diversification during phage predation and human infection using sensitive NGS IS-Seq. This work provides a high-resolution picture of IS256 movement during both steady-state and stress inducing conditions. Understanding the outcomes of these IS256 insertion events will be critical to the study of the evolution of enterococcal pathogenesis in response to both phage and antibiotic therapies. We also show that IS256 is regulated at the transcriptional level in *E. faecalis* and have discovered that IS256 mobility is biased for highly mobilizable insertion events in nosocomial and virulent enterococcal isolates, further demonstrating that IS elements are domesticated and specialized in their choice of bacterial host.

## Materials and Methods

### Bacteria and bacteriophages

*E. faecalis* strains were grown in Brain Heart Infusion (BHI) broth with aeration at 37°C. A list of bacterial and bacteriophage strains can be found in Table S4. *Escherichia coli* cultures were grown in Lennox L broth at 37°C with aeration. 15 μg/mL or 5 μg/mL chloramphenicol was used for selection of transformed *E. coli* and *E. faecalis*, respectively. The generation of phage-resistant 19RS strains was recently described [18].

### Enterococcal phylogeny and Identification of IS256 in enterococcal genomes

All available *E. faecalis* and *E. faecium* genomes were downloaded from NCBI RefSeq on 3/21/2022. Phylogenetic trees were constructed with GToTree v1.6.31 using default parameters [96], pruned using treeshrink v1.3.9 [97], and visualized using the iTOL viewer [98]. IS256 transposase genes were identified using GToTree with the IS256 pfam “PF00872” and were further refined by selecting matches with 100% amino acid matches to the IS256 Tnp from *E. faecalis* V583 using seqkit v0.16.1 [99].

### MLST and virulence factor abundance

*Enterococcal* genomes obtained from RefSeq were assigned an MLST type using mlst v2.19.0 [100] with the *E. faecalis* and *E. faecium* MLST scheme from pubmlst [101]. Virulence factors were identified by calling all predicted ORFs in all downloaded RefSeq genomes using prodigal v2.6.3 [102] and running diamond blastp v0.9.21 [103] against this database with known *E. faecalis* and *E. faecium* virulence factors from the Virulence Factor Database [104]. All further analyses and statistics were performed in R.

### DNA extraction

Bacterial genomic DNA was extracted from overnight cultures grown in 5 mL BHI broth using the Zymo BIOmics DNA Miniprep Kit (Zymo, Irvine, CA).

### Whole genome sequencing, IS256 identification from whole genome sequencing, and sequencing read alignment to the phi19 genome

Whole genome sequencing of *E. faecalis* V583 and 19RS strains was performed using Illumina HiSeq 2500 with 150 cycle paired-end sequencing. These genomes were previously described [18], and the reads have been deposited at the European Nucleotide Archive (ENA) as project accession number PRJEB30526. Novel IS256 insertions were found using CLC Workbench (Qiagen, Hilden, Germany) by first mapping reads to an *E. faecalis* V583 reference genome where all IS256 elements were removed manually. Unmapped reads were collected and mapped against a database of known IS256 elements with mapping parameters that allowed for partial matches (0.4 length fraction and 0.9 fraction similarity). Reads that mapped against IS256 were binned, the section of the read containing the IS256 sequence was removed, and the remaining read was realigned against the *E. faecalis* V583 reference genome (GCA_000007785.1) to identify IS256 insertion sites.

Reads aligning to the phi19 genome were found by first aligning whole genome sequencing reads to the *E. faecalis* V583 reference genome using Bowtie2 v2.3.5.1[105], collecting unmapped reads, and aligning these reads to the phi19 genome.

### Oxford Nanopore Technologies (ONT) sequencing

High molecular weight genomic DNA was isolated from *E. faecalis* V583 and 19RS strains after overnight growth as described above. ONT sequencing was performed by SeqCenter (Pittsburgh, PA) using PCR-free V14 chemistry ligation library preparation and sequenced on a GridION (ONT, Oxford, UK). 113,000 ± 22,400 reads were generated with an average read length of 3.7 ± 0.8 kb . phi19 sequences in reads were found using blastn [106] with a minimum alignment length of 1 kb. Reads passing this threshold were compared to the *E. faecalis* V583 genome with blastn and positive hits were filtered with a minimum alignment length of 1 kb. The reads have been deposited at the ENA as project accession number PRJEB30526.

### IS-Seq analysis

For library preparation, 200 ng of genomic DNA was used as input in the Illumina DNA Prep Kit following the manufacturer’s instructions (Illumina, San Diego, CA). For the in vitro and in vivo experiments (Fig. 4), genomic DNA was isolated from 5x10^9^ *E. faecalis* cells. Following library construction, IS256 enrichment was performed as follows. First, 10 ng of library DNA was amplified with the p7 primer and the IS-Seq Step1 primer (Table S5) for 13 cycles using Q5 2X Master Mix (New England Biolabs, Ipswich, MA). These reaction products were diluted 1:100 and 10 µL was added to a PCR reaction with the p7 primer and the IS-Seq Step2 primer for 9 cycles using Q5 2X Master Mix. The products of this second reaction were sequenced on an Illumina NovaSEQ 6000 by the University of Colorado Anschutz Medical Center Genomics Core with paired-end 150 cycle chemistry. 23.3 ± 3.8 M reads, 17.9 ± 2.5 M reads, and 16.3 ± 2.6 M reads were obtained from sequencing the human, mouse, and *in vitro* IS-Seq libraries, respectively. IS-seq reads have been deposited at the ENA under project accession number PRJEB55280.

Paired end reads obtained from IS-Seq were partitioned into read1 and read2 files, and reads from read1 files were used in the downstream analysis. First, cutadapt v1.18 was used to bin reads with a full IS256 5’ terminus. The IS256 5’ terminus was then trimmed from these reads using cutadapt with the following flags: ‘-O 6 -g "^CGTAAAAGGACTGTTATATGGCCTTTTTACTTTTACACAATTATACGGACTTTATC" ‘ [107]. These trimmed reads were aligned to the *E. faecalis* V583 reference genome using Bowtie2 and the location of each insertion was determined using samtools v1.13 [108] and bedtools genomecov v2.30.0 [109].

For data generated in Figure 7, contigs were co-assembled from Illumina sequencing reads from each sample using megahit v. 1.2.7 [110]. These Illumina reads were generated from libraries prepared using the Nextera prep kit (Illumina) and sequenced on an Illumina NextSeq 500 with 150 cycle paired-end sequencing. Duplicate contigs were removed using gclust v1.0 [111] and these representative contigs were used as a reference for aligning IS-Seq reads as described above.

All further downstream analysis was performed in R. Insertion site abundances were normalized using DESeq2 estimateSizeFactors [56] before plotting to account for abnormalities during library preparation and sequencing. Significantly enriched unique insertion sites were found using DESeq2 by providing raw counts to the program and then computing general linearized models for each insertion site with at least five reads in three replicates. Models were fit using either a negative binomial fit, or when appropriate, a local fit. To account for fold change variance during each pairwise comparison, log fold change values were shrunk using the ApeGLM model [112] in DESeq2’s *lfcshrink* function. Each pairwise comparison was performed using a separate call to the *DESeq* function in order to improve sensitivity, as recommended by the developers [56]. Both significantly enriched and insignificant insertions with at least 100 reads were reported in tables S2, S3 and figures 4B, 4D, 7B.

### Stranded RNA-Seq analysis

The *E. faecalis* V583 RNA-Seq data used for stranded IS256 gene regulation analysis was obtained from SRA experiments SRR7229491, SRR7229489, and SRR7229487 [54]. The reads were aligned to an *E. faecalis* V583 IS256 element sequence using Bowtie2 and forward and reverse aligning reads were found by parsing the SAM file. Only reads that perfectly aligned to the IS256 reference were retained. The depth of read alignment at each position in IS256 was calculated using samtools *depth* and the data was visualized in R.

### E. faecalis phi19 infection in the mouse intestine

All animal protocols were approved by the Institutional Animal Care and Use Committee of the University of Colorado School of Medicine (protocol number 00253). Conventional 6-week-old C57BL/6 mice were divided into two groups (Control: 2 female/ 2 male, Experimental: 2 female/ 2 male). All mice were treated with an antibiotic cocktail (streptomycin [1Lmg/ml], gentamicin [1Lmg/ml], erythromycin [200Lμg/ml]) by first dosing via oral gavage with 100 µl of the cocktail and replacing their drinking water with the same antibiotic cocktail for 1 week. On day 7, antibiotic water was replaced with antibiotic free-drinking water. After 24 hours mice were colonized with *E. faecalis* V583 suspended in phosphate buffered saline (PBS) by oral gavage (10^9^ CFU in 100 µl). After 24 hours mice were orally gavaged daily as follows: 100 μl of 1M sodium bicarbonate (all mice) and 100 µl PBS per control mouse or 10^9^ PFU phi19 in 100 µl per experimental mouse. Feces were collected daily and homogenized in 1ml PBS. 10µl of the fecal slurry was serially diluted and plated on BHI agar containing 100 μg/ml gentamicin or 100 μg/ml gentamicin and 10^8^ pfu/ml phi19. From the experimental samples, 5 µl of slurry was mixed with 45µl chloroform, spun at 16,363 RCF for 1 min and the supernatant was enumerated for phage particles by agar overlay plaque assay as described previously [19]. *E. faecalis* colonies were isolated on BHI agar containing 100 μg/ml gentamicin. These cells were grown to stationary phase in 5 mL of BHI broth, after which genomic DNA was isolated and IS-Seq was performed on these samples.

### RNA extraction and qRT-PCR

RNA was isolated from exponentially growing (OD=0.3) *E. faecalis* using a modified protocol for the RNeasy Mini-Prep Kit (Qiagen). First, cell pellets were collected by centrifugation of 5 mL of cell culture at 8,228 RCF for 5 minutes. Cell pellets were resuspended in 1 mL RNAlater and centrifuged again at 8,228 RCF for 10 minutes. The cell pellets were stored at -80 °C until sample processing. To isolate RNA, the pellets were thawed and resuspended in 100 μL TE buffer with 15 mg/mL lysozyme and incubated at room temperature for 30 minutes. Following this, 700 μL Buffer RLT with 0.01% beta-mercaptoethanol was added and the sample was transferred to a Lysing Matrix B tube (MPBio, Irvine, CA). The tubes were bead-beaten in a Mini-Beadbeater-16 (Biospec Products, Bartlesville, OK) on the fastest setting at 30-second intervals with 1-minute rests on ice between cycles. Following bead beating, the supernatant was removed after centrifugation at 16,363 RCF for 30 seconds and processed following the manufacturer’s instructions. RNA was eluted from the column in RNase/DNase free H_2_O and any residual DNA contamination was removed with a 1 hour off-column DNase treatment (Qiagen). The RNA was purified following the RNeasy Mini-Prep Kit following manufacturer’s instructions. cDNA was synthesized using Qscript Master Mix (Quantabio, Beverley, MA) and 1 μg RNA. For qPCR of gDNA targets, 1 ng of gDNA template was used per reaction. qPCR was run with PowerUp SYBR Green Master Mix (Thermo Fisher, Waltham, MA) on an Applied Biosystems QuantStudio 7 Flex qRT-PCR system. qRT-PCR primers are listed in Table S5. Total 16S rRNA gene transcripts were used for qRT-PCR normalization and *clpX* was used for DNA qPCR normalization associated with IS circle quantification.

### DNA manipulation and cloning

All cloning primers and DNA constructs used in this study are listed in Table S5. All plasmid constructs used the pLZ12 shuttle vector backbone [113]. For cloning, all inserts and vectors were amplified using 2X Q5 DNA polymerase Master Mix and assembled using 2X Gibson Assembly Master Mix (New England Biolabs).

### Phage carriage identification and serial passaging

To enumerate extracellular viral particles, we collected supernatant from cultures after centrifugation for 1 minute at 10,000 RCF, treated with 1/10 volume chloroform, centrifuged for 1 min at 21,000 RCF and extracted the aqueous phase. The resulting supernatant was serially diluted in SM+ buffer (50 mM Tris-HCl, 100 mM NaCl, 8LmM MgSO_4_, 5LmM CaCl_2_ [pH 7.4]), and 10 μL was mixed with 130 μL of a 1:10 dilution of *E. faecalis* V583 cells grown overnight. This mixture was combined with 5 mL of 0.35% Todd Hewitt agar (THA) supplemented with 10mM MgSO_4_, and poured onto THA+10mM MgSO_4_ plates.

To identify 19RS *E. faecalis* cells actively shedding phages, glycerol stocks of phage-resistant cultures were streaked on BHI agar and incubated at 37°C. After overnight growth, 182 μL of a 1:10 dilution in SM+ of an overnight wild type *E. faecalis* V583 culture was mixed with 7 mL of 0.35% THA and 10mM MgSO_4_ and solidified on THA+10mM MgSO_4_ plates for 30 minutes. Individual colonies were picked from the overnight plate patched onto a plate containing *E. faecalis* V583 embedded in 0.35% THA, with care not to break the surface of the agar, and incubated at 37°C overnight.

To serially passage phage-positive colonies, colonies producing zones of clearing were patched onto both THA+10mM MgSO_4_ plates and THA+10mM MgSO_4_ with an *E. faecalis* V583 top agar layer as described above. The next day, the colony on the THA+10mM MgSO_4_ plate was patched onto both types of plates and this was continued for two subsequent passages.

### Competition assays

*E. faecalis* cultures were grown overnight in BHI broth and diluted to an OD_600_ of 1 in PBS. 30 μL of each strain was added to 3 mL BHI and incubated at 37°C for 24 hours. The cultures were plated on both BHI gentamicin 100 μg/mL to select for *E. faecalis* V583 strains and BHI rifampicin 25 μg/mL, fusidic acid 50 μg/mL to select for *E. faecalis* OG1RF or *E. faecalis* OG1RF Δ*epaOX*. Competitive fitness was calculated by dividing the final concentration of *E. faecalis* OG1RF or *E. faecalis* OG1RF Δ*epaOX* by the final concentration of *E. faecalis* V583 or *E. faecalis* 19RS and logarithmically scaling this quotient.

### Growth curves

*E. faecalis* cultures were grown overnight in BHI broth and diluted to an OD_600_ of 1 in PBS. 50 μL of each strain was added to 50 mL BHI and incubated at 37°C. OD_600_ readings were taken every 30 minutes on a Tecan Infinite M Plex (Tecan, Männedorf, Switzerland).

### Enterococcal isolation from stool samples and gDNA extraction

Enterococcal stool populations were obtained over a span of 109 days from a patient being treated for recurrent *E. faecium* bacteremia. Stool samples were diluted and plated onto bile esculin azide – Enterococcosel agar (Becton Dickinson, Franklin Lakes, NJ), and 100-1,000 colonies were collected and pooled into BHI containing 16.7% glycerol and stored at -80°C. One mL aliquots of the frozen population stocks were thawed and washed with BHI. Populations were allowed to briefly expand at 37°C at 170 rpm for 5.5 hours. The cultures were pelleted and genomic DNA was extracted into DNase free water using the DNeasy Blood and Tissue Kit (Qiagen). gDNA concentrations were determined with the Qubit 1X dsDNA High-Sensitivity Kit and fluorometer (Invitrogen, Thermo Fisher).

### Phage analysis from clinical enterococcal blood isolates

Viral contigs were found in assembled contigs using VIBRANT v1.2.1 [114] and reads from clinical samples were mapped against these contigs using Bowtie2. Multiple comparison analysis was performed using DESeq2 as described above.

### Statistics and Visualization

Graphs were made in Graphpad Prism or in R using dplyr and ggplot2. t-tests and one-way ANOVAs were calculated in Graphpad Prism. Chi-squared tests and insertion site differential abundance and proportional tests (prop.test) were calculated in R. Significantly abundant insertions from the IS-Seq experiments were found with DESeq as described above.

## Acknowledgements

This work was supported by National Institutes of Health grants R01AI141479 (B.A.D.), R01AI165519 (D.V.T.), R01AI116610 (K.L.P.), T32AI052066 (J.M.K), F31AI169976 (J.M.K.), and T32AI138954 (M.E.S.).

## Supplementary Figure Legends

**Figure S1. Enrichment of IS256-chromosomal junctions during IS-Seq. A)** Schematic of the IS-Seq PCR amplification step. **B)** Trimming site and binning strategy used to identify IS256-termini reads.

**Figure S2. Steady state IS256 insertion sites in the native *E. faecalis* V583 plasmids. A)** pTEF1, **B)** pTEF2, and **C)** pTEF3. Three individual biological replicates are shown in each panel.

**Figure S3. Diagram of IS256 mutant constructs.**

**Figure S4. IS256 insertions in *E. faecalis* V583 plasmids in vitro and in vivo. A-C)** Insertion sties from in vitro cultured *E. faecalis* 19RS strains; **A)** pTEF1, **B)** pTEF2, and **C)** pTEF3. **D-F)** Insertion sites from *E. faecalis* isolated from the murine intestine **D)** pTEF1, **E)** pTEF2, and **F)** pTEF3. Each peak in panels A, B, and C are the average of three biological replicates per strain, and in panels D, E, and F are an average of two biological replicates.

**Figure S5. IS-256 insertions in *epa* and *vex*/*vnc* genes in vitro and in vivo.** Insertion sties from in vitro cultured *E. faecalis* 19RS strains in the **A)** *epa* locus and **B)** *vex*/*vnc* locus. Insertion sites from *E. faecalis* isolated from the murine intestine in the **C)** *epa* locus and **D)** *vex*/*vnc* locus. Each peak in panels A and B are the average of three biological replicates per strain, and in panels C and D are an average of two biological replicates.

**Figure S6. Murine model and intestinal *E. faecalis* colonization and oral phi19 treatment. A)** Schematic of the *E. faecalis* intestinal colonization model used to identify in vivo IS256 mobilization. The bacterial population was isolated during the phage treatment phase (between Day 9-22) by plating on selective media and genomic DNA was isolated from these cells. **B)** Colony forming units of *E. faecalis* isolated from mouse feces. Bacteria were enumerated on growth media supplemented with gentamicin with and without the addition of phi19 to measure the frequency of phage resistant colonies. Arrows indicate time points where IS-Seq was performed. Four mice were used in both the phi19 treated and untreated groups. Bacteria used for IS-Seq characterization were isolated on BHI gentamicin media.

**Figure S7. Serial passage of phi19 shedding *E. faecalis* 19RS isolates.**

**Figure S8. IS256 read mapping to enterococcal genomic contigs assembled from stool samples.** Each data point indicates the read mapping abundance per assembled contig. Data points indicated as *E. faecalis* or *E. faecium* lacked sufficient data to definitively categorize these contigs as either *E. faecalis* or *E. faecium*.

**Figure S9. IS256 insertions from human stool enterococcal isolates in genes involved in aminoglycoside resistance.** Each insertion is derived from one biological replicate per time point from the patient’s stool.

**Figure S10. Differential abundance of predicted prophages in clinical samples.**

## Supplementary Table Legends

**Table S1. MLST abundances and virulence factor abundances in IS256 and non-IS256 E. faecalis and E. faecium genomes.**

**Table S2. Read counts and differential abundance analysis of abundant IS256 insertions in E. faecalis V583 and 19RS strains.**

**Table S3. Read counts and differential abundance analysis of abundant IS256 insertions in *E. faecalis* V583 isolated from mouse stool.**

**Table S4. List of bacteria and bacteriophages used in this study.**

**Table S5. List of primers and plasmids used in this study.**

